# Sensitivity of infectious SARS-CoV-2 B.1.1.7 and B.1.351 variants to neutralizing antibodies

**DOI:** 10.1101/2021.02.12.430472

**Authors:** Delphine Planas, Timothée Bruel, Ludivine Grzelak, Florence Guivel-Benhassine, Isabelle Staropoli, Françoise Porrot, Cyril Planchais, Julian Buchrieser, Maaran Michael Rajah, Elodie Bishop, Mélanie Albert, Flora Donati, Sylvie Behillil, Vincent Enouf, Marianne Maquart, Maria Gonzalez, Jérôme De Sèze, Hélène Péré, David Veyer, Aymeric Sève, Etienne Simon-Lorière, Samira Fafi-Kremer, Karl Stefic, Hugo Mouquet, Laurent Hocqueloux, Sylvie van der Werf, Thierry Prazuck, Olivier Schwartz

## Abstract

SARS-CoV-2 B.1.1.7 and B.1.351 variants emerged respectively in United Kingdom and South Africa and spread in many countries. Here, we isolated infectious B.1.1.7 and B.1.351 strains and examined their sensitivity to anti-SARS-CoV-2 antibodies present in sera and nasal swabs, in comparison with a D614G reference virus. We established a novel rapid neutralization assay, based on reporter cells that become GFP+ after overnight infection. B.1.1.7 was neutralized by 79/83 sera from convalescent patients collected up to 9 months post symptoms, almost similar to D614G. There was a mean 6-fold reduction in titers and even loss of activity against B.1.351 in 40% of convalescent sera after 9 months. Early sera from 19 vaccinated individuals were almost as potent against B.1.1.7 but less efficacious against B.1.351, when compared to D614G. Nasal swabs from vaccine recipients were not neutralizing, except in individuals who were diagnosed COVID-19+ before vaccination. Thus, faster-spreading variants acquired a partial resistance to humoral immunity generated by natural infection or vaccination, mostly visible in individuals with low antibody levels.

---

SARS-CoV-2 variants rapidly emerged in humans and supplanted ancestral strains^1–5^. Their increased rates of inter-individual transmissions conferred a replication advantage at the population level. One of the first identified variant includes the D614G mutation in the Spike protein, which enhances viral infectivity and shifts Spike conformation towards an ACE2-binding fusion-competent state, without significantly modifying sensitivity to antibody neutralization ^1,6–8^. More recently, novel variants appeared in different countries, with combinations of mutations and deletions in the RBD and N-terminal Domains (NTD) of Spike as well as in other proteins. The B.1.1.7 variant emerged in the UK, the B.1.351 variant (also termed 501Y.V2) in South Africa and the P.1 and P.2 lineages in Brazil ^2,3,5,9–12^. Although different, the variants share common characteristics, including known escape mutations that were previously identified under antibody pressure selection in vitro ^2,3,13–17^. Some of the mutations or deletions were also identified in immuno-compromised individuals with prolonged infectious viral shedding and treated with convalescent plasma or anti-Spike monoclonal antibodies ^3,17–19^, indicating that antibody escape mutations are selected in vivo. The sensitivity to antibody neutralisation varies with the viral variant. B.1.1.7 seems to be more sensitive to neutralization than B.1.351. The RBD mutation N501Y that increases affinity to ACE2 and is present in B.1.1.7 and B.1.351 ^20^ does not impair on its own post-vaccine serum neutralization ^21^. The other mutations in B.1.1.7 have been suggested to not result in immune evasion of linear epitopes ^22^. Mutations in B.1.351 and P.1 strains, including E484K and K417N/T, are of high concern, since they partly compromise neutralization generated by previous infection or vaccination ^23–25^ or may increase inherent viral fitness. The Pfizer Cominarty™ (also termed BNT162b2) vaccine–elicited human sera neutralizes SARS-CoV-2 lineage B.1.1.7 pseudovirus, with slightly reduced titers in some vaccinees, when compared to the Wuhan reference strain ^26,27^. The Moderna mRNA-1273 vaccine also induces neutralizing antibodies against SARS-CoV-2 variants, with however a 5-10 fold reduction in efficacy against the B.1.351 Spike, when compared to pseudovirus bearing the D614G mutation ^27,28^.

Neutralization efficacy was so far mostly assessed using VSV-derived or lentivirus-derived pseudovirus assays, or with infectious SARS-CoV-2 carrying point mutations in the Spike. One recent report using infectious B.1.351 virus showed that plasma from six convalescent donors were strongly attenuated against this strain, with 6 to 200-fold higher IC50 values relative to ancestral virus ^29^. It is thus of outmost importance to use authentic variant strains in addition to pseudovirus particles to evaluate their sensitivity to antibodies. Here, we compared the sensitivity of three authentic SARS-CoV-2 strains, the pre-existing D614G virus and the B.1.1.7 and B.1.351 variants, to antibody neutralization.

## Results

### Isolation and characterization of SARS-CoV-2 B.1.1.7 and B.1.351 variants

We isolated the two variants from nasal swabs of patients with RT-qPCR and sequence-diagnosed infection. The viruses were amplified by one to two passages on Vero cells. Sequences of the outgrown viruses confirmed the identity of B.1.1.7 and B.1.351 variants. Viral stocks were titrated using S-Fuse reporter cells ^30^. These cells are derived from U2OS-ACE2+ cells and carry the GFP-split system. Upon infection, the cells produce the Spike at their surface and fuse with neighbouring cells, generating a GFP signal after a few hours ^30^ (Fig. 1a). The number of GFP+ cells correlated with the viral inoculum (Fig. 1b). Neutralizing anti-SARS-CoV-2 monoclonal antibodies (mAb) targeting the RBD can be classified into 4 main categories ^31,32^. We tested with the S-Fuse assay the sensitivity of the three viral strains to two anti-RBD antibodies, mAb102 and mAb48, that belong to the first category, and act by blocking ACE2 and binding to “up” RBDs ^32^. MAb102 efficiently and similarly neutralized the three viral strains, with IC50 of ≃ 0.01 μg/mL (Fig. 1c,d). MAb48 neutralized D614G (IC50 of 0.1 μg/mL) but was inactive against B.1.1.7 and B.1.351 variants (Fig. 1c,d). These results confirmed that the two variants selectively display reduced sensitivity to certain antibodies. Of note, the activity of the mAbs was similar at different viral inocula, within a range of 50 to 200 GFP+ syncytia/well (Fig. 1e), indicating that potential variations in the number of infected S-Fuse cells do not impact the calculation of IC50. We thus selected for further studies a multiplicity of infection leading to about 150 syncytia/well for each viral strain.

**Fig. 1.**
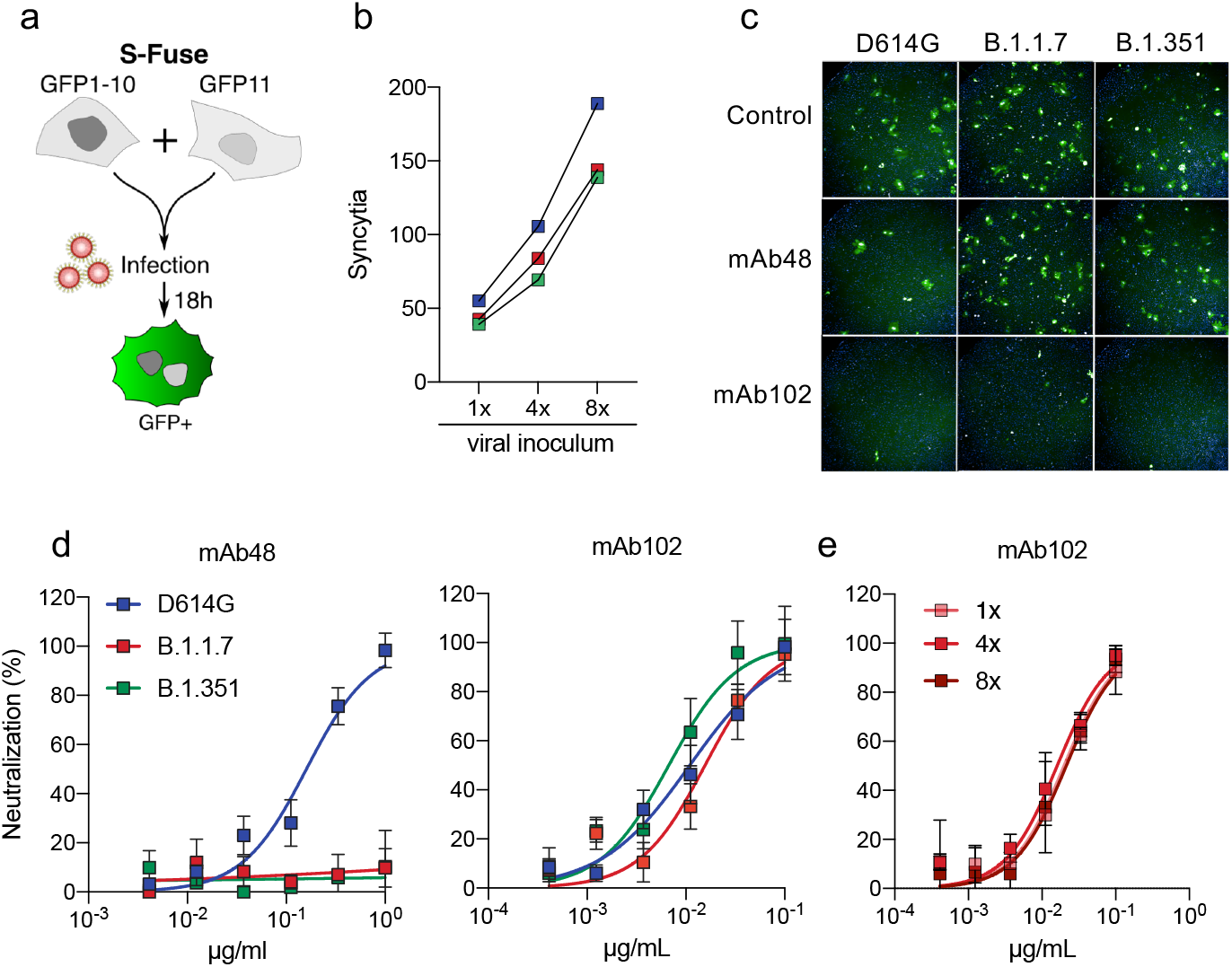
Effect of two anti-SARS-Cov-2 monoclonal antibodies on D614G, B.1.1.7 and B1.351 strains. **a.** Principle of the S-Fuse reporter assay. S-Fuse cells, a mix of U20S-ACE2-GFP1-10 and U20S-ACE2-GFP11 were cocultured at a 1:1 ratio and infected with SARS-CoV-2. Cells fuse upon infection and generate a GFP signal. Infection was quantified by measuring the number of GFP+ syncytia at 18 hours. **b.** Linearity of the assay with D614G (blue), B.1.1.7 (red) and B.1.351 (green) strains. S-Fuse cells were exposed to serial dilutions of viral stocks and infection was quantified by high content imaging. **c.** Representative images of S-Fuse cells infected with D614G, B.1.1.7 and B1.351 strains in the presence or absence of two anti-SARS-CoV-2 monoclonal antibodies (mAb48 and mAb102). **d.** Dose response analysis of the neutralization by mAb102 and mAb48 on the three viral strains. **e.** Neutralization of SARS-CoV-2 (B.1.1.7) by mAb102 at three viral inocula. Results are shown as mean ± SD from four independent experiments.

### Sensitivity of D614G, B.1.1.7 and B.1.351 variants to sera from convalescent individuals

We then assessed the neutralization ability of sera from convalescent individuals. We randomly selected 28 samples in a longitudinal cohort of infected individuals from the French city of Orléans (Table S1). They were all diagnosed for COVID-19 by RT-qPCR or serology and included critical, severe, and mild-to-moderate cases. 25 individuals were sampled twice, firstly at a median of 89 days (range: 77-105) post onset of symptoms (POS) (Month 3 samples (M3)), and secondly at a median of 179 days (range 168-197) POS (M6 samples). We incubated serially diluted sera with D614G, B.1.1.7 or B.1.351 strains, added the mixture to S-Fuse cells, and scored the GFP+ cells after overnight infection. We then calculated ED50 (Effective Dose 50%) for each combination of serum and virus. A representative example with the same donor is depicted Fig. 2a, and the results with all donors appear in Fig. 2b. The D614G and B.1.1.7 strains were similarly sensitive to sera. At M3, the median ED50 values were respectively 1.5×10^3^ and 1×10^3^ for D614G and B.1.1.7 variants, with huge variations between individuals (from 10^2^ to 2×10^4^) and the values did not strongly decline at M6. As expected, individuals with critical conditions displayed higher neutralizing activities than those with severe or mild-to-moderate symptoms (Fig. S1). Again, there was no significant difference between the two viral strains in each category of symptoms. With B1.351, the neutralization titers were significantly decreased by 5 and 10-fold at the two time-points, when compared with D614G and B.1.1.7 strains (Fig. 2b and S1). We confirmed these results in 30 sera from another cohort of RT-qPCR-confirmed staff from Strasbourg University Hospitals that experienced a mild disease ^33,34^. The samples were collected at a later time point (M9), with a median of 233 days (range 206-258) POS (Table S1). Overall, the neutralization activity was lower (one representative example is shown in Fig. 2a and the results of all donors in Fig. 2b). There was no significant difference in neutralization between D614G and B.1.1.7 strains, with a similar ED50 of 2×10^2^. In sharp contrast, the neutralizing activity against B.1.351 was particularly low at this late time point, with a median ED50 of 50, representing a 4-fold decrease when compared to D614G (Fig. 2b).

**Fig. 2.**
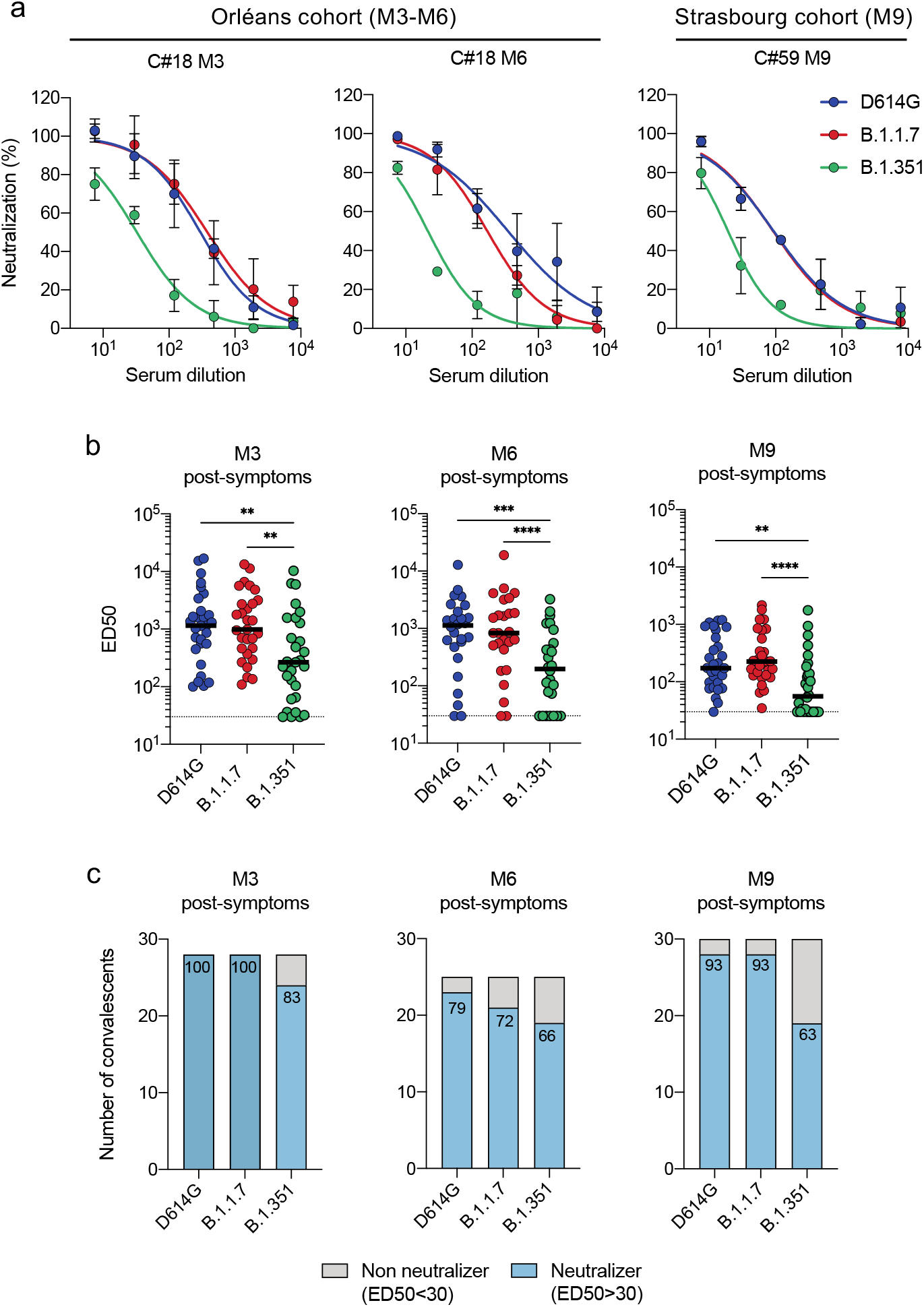
Sensitivity of SARS-CoV-2 D614G, B.1.1.7 and B.1.351 variants to sera from convalescent individuals. **a.** Examples of neutralization curves with sera from two donors. The first donor, from the Orléans Cohort, was sequentially sampled at month 3 (M3) and M6 (left and middle panels, respectively). The second donor, from the Strasbourg Cohort, was sampled at M9 (right panel). Results are shown as mean ± SD from two to four independent experiments. **b.** ED50 of neutralization of the three viral isolates. 28 sera from the Orléans Cohort were sequentially sampled at month 3 (M3) and M6 (left and middle panels, respectively), and 30 sera from the Strasbourg Cohort were sampled at M9 (right panel). Data are mean from two to four independent experiments. Friedman test with Dunn’s multiple comparison was performed between each viral strain at the different time points, *: p-value<0.05 **: p-value<0.01 **** p<0.001. **c.** Each subject was arbitrarily defined as “neutralizer” (in blue) if a neutralizing activity was detected at the first (1/30) serum dilution or “non-neutralizer” (in grey) if no activity was detected. The numbers indicate the % of neutralizers.

We then arbitrarily classified the cases as neutralizers (with neutralizing antibodies detectable at the first serum dilution of 1/30) and non-neutralizers, for the three viral strains and the two cohorts (Fig. 2c). Most of individuals neutralized the three strains at M1. The fraction of neutralizers started to decline at M6, a phenomenon which was more marked with B.1.351. The fraction of neutralizers was higher in the second cohort, with 93% of individuals neutralizing either D614G or B.1.1.7 strains at M9. This fraction however dropped to 63% of neutralizers against the B.1.351 strain (Fig. 2c).

### Antibody binding to cells expressing D614G, B.1.1.7 and B.1.351 Spikes

We then examined the binding capacity of monoclonal antibodies and sera to Spikes from different lineages. To this aim, we adapted the flow cytometry based S-Flow assay, that we previously established to measure the levels of antibodies binding to cells expressing the Wuhan Spike ^35^. We transiently transfected 293T cells with plasmids expressing the D614G, B.1.1.7 and B.1.351 Spikes. Similar surface levels of the viral proteins were detected with mAb10, a non-neutralizing pan-coronavirus antibody targeting a conserved epitope in the S2 domain (Fig. 3a,b and Fig. S2). Moreover, the cells fused when co-cultivated with ACE2-expressing cells, indicating that the three proteins were functional (not shown). The mAb102 similarly bound to the three Spikes, in accordance with its cross-reactive neutralizing activity. mAb48 efficiently bound to D614G Spikes, slightly less potently to B.1.1.7, and lost any binding activity to B.1.351. Therefore, the escape of mAb48 neutralization by the two variants is due to mutations decreasing or abrogating antibody binding to its target. The N501Y mutation enhances RBD affinity to ACE2, when tested with recombinant proteins and yeast surface display ^20^. We assessed by flow cytometry the binding of a labelled soluble ACE2 protein to cells expressing the different Spikes. We observed a dramatic increase of binding of soluble ACE2 to B.1.1.7 and to a lesser extent to B.1.351, which both carry the N501Y mutation, when compared to D614G (Fig. 3a,b).

**Fig. 3.**
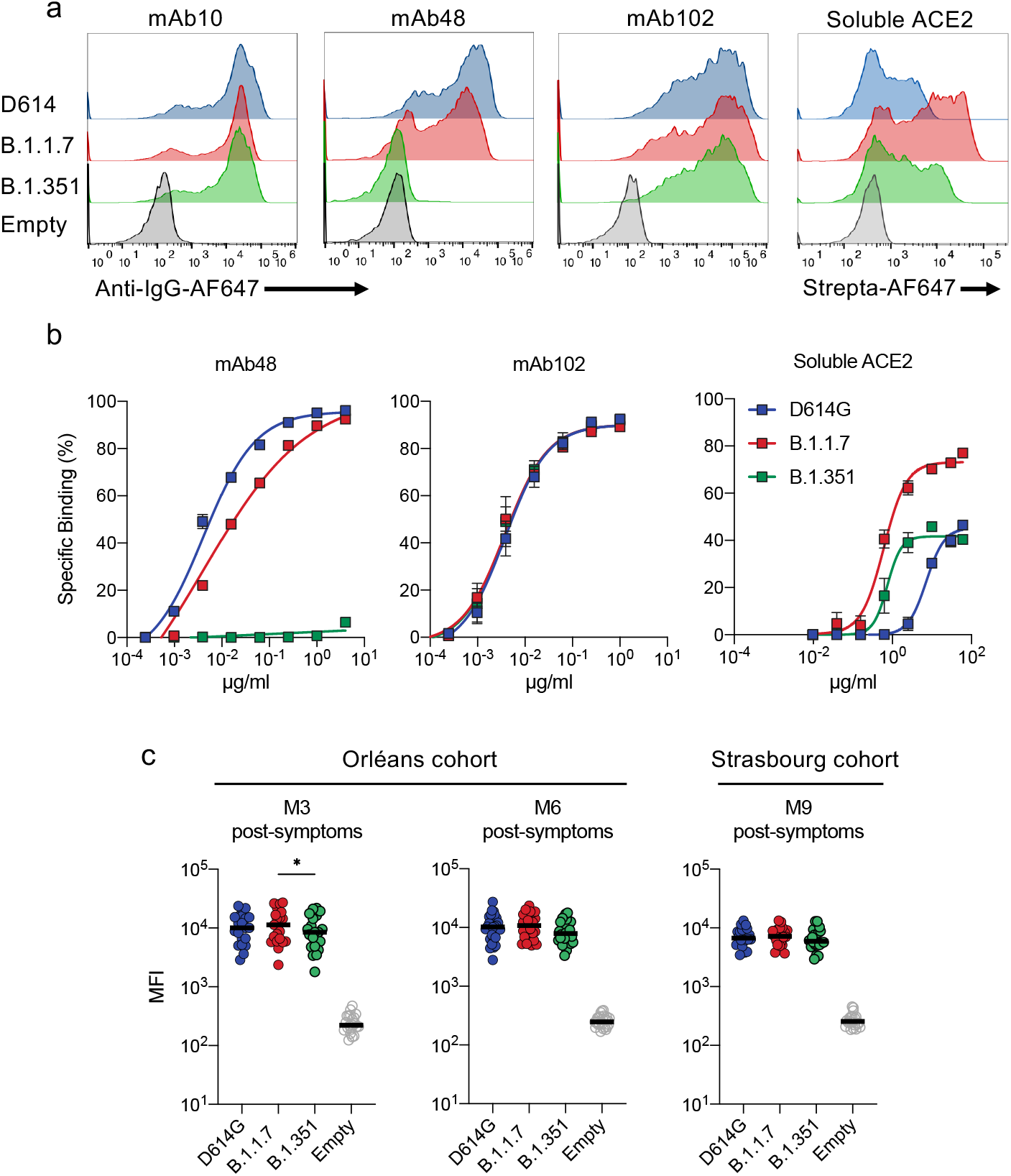
Antibody binding to cells expressing D614G, B.1.1.7 and B.1.351 Spikes. **a.** Binding of monoclonal antibodies or soluble ACE2. 293T cells were transiently transfected with plasmids expressing the D614G, B.1.1.7 and B.1.351 Spikes. After 24h, cells were stained with anti SARS-CoV-2 antibodies mAb10 (a pan-coronavirus antibody), mAb48, mAb102 or soluble ACE2 (ACE2-biotin at 10 μg/ml revealed with fluorescent Streptavidin) and analyzed by flow-cytometry. One representative example of binding is shown. **b.** Titration binding of curves of mAb48, mAb102 and ACE2 to the three Spikes. Data are mean of three independent experiments. **c,d.** Binding of the panel of 83 sera from 58 convalescent individuals. Sera were tested at a 1/300 dilution. Data are mean of two independent experiments.

We next tested our panel of 83 sera from convalescent individuals against the different Spikes. Overall, the sera similarly bound to the Spikes, even though B.1.351 displayed a slight reduction in the mean fluorescence intensity (MFI) of binding (Fig. 3c). We observed a global and slight decrease of MFI at M9, compared to samples from M3 and M6 (Fig. 3d). Altogether, these results indicate that Spikes from B.1.351 and to an higher extent B.1.1.7 display increased affinity for ACE2, while escaping binding to some monoclonal antibodies and either to a lesser degree or not at all to polyclonal sera.

### Sensitivity of D614G, B.1.1.7 and B.1.351 variants to sera from vaccine recipients

We next asked whether vaccine-elicited antibodies inhibit infection by the different variants. In France, vaccination started in January 2020 with the Pfizer Comirnaty vaccine. We thus selected 19 vaccine recipients from a cohort of vaccinated health care workers established in Orléans. The characteristics of vaccinees are depicted Table S2. Sera and nasal swabs were sampled at week 2 (17 individuals) and week 3 (18 individuals), week 4 (11 individuals, corresponding to week 1 after the second dose), with a median of 13, 19 and 28 days after the first dose, respectively. This allowed us to assess the early humoral response to vaccination. We first analyzed the 16 out of 19 vaccinees that have not been previously infected with SARS-CoV-2, as assessed by the absence of pre-existing anti-S antibodies and anti-N antibodies. A representative example of the evolution of the neutralizing response in one vaccine recipient at the three time-points is depicted Fig. 4a. In this case, the serum only neutralized D614G at week 2, whereas B.1.1.7 started to be neutralized at week 3, although less efficiently than D614G. B.1.1.7 and D614G strains were similarly neutralized at week 4. The anti-B.1.351 response was negative up to week 3, and became detectable at week 4, at a level lower than the two other viruses. The ED50 results with the 16 vaccine recipients are presented Fig. 4b. We also arbitrarily classified the cases as neutralizers (with neutralizing antibodies detectable at the first serum dilution of 1/30) and non-neutralizers, for the three viral strains (Fig. 4c). Two weeks after vaccination, antibodies neutralizing the D614G strain were detected in the sera of 5/15 recipients (33%, with an arbitrary threshold of ED50> 30 for neutralization positivity) (Fig. 4b,c). The titers were relatively modest at this early time point (median ED50: 30). These low titers were less efficient against B.1.1.7, with 2/15 neutralizers (13%), and inactive against B.1.351 (Fig. 4b,c). At week 3, the neutralizing activity increased against D614G and B.1.1.7. There was however a 3-fold reduction in the neutralization titers with B.1.1.7 (Fig. 4b), whereas B.1.351 remained unsensitive. At the later time point (week 1 after the second dose), titers increased in most of the recipients, were similar between D614G and B.1.1.7, but remained 7-fold lower with B.1.351 (Fig. 4b). At week 3, 63%, 38% and 0% neutralized D614G, B.1.1.7 and B.1.351 strains, respectively (Fig. 4c). These percentages increased after the second dose, to reach 80% of neutralizers for D614G and B.1.1.7, whereas 60% neutralized B.1.351 with a low titer (Fig. 4c). The S-Flow demonstrated the presence of antibodies binding to the three Spikes at the three sampling times, with a noticeable decrease for the B.1.351 Spike (Fig. 5). Therefore, the Pfizer vaccine generated a neutralizing response that efficiently targeted D614G and B.1.1.7. There was a delay in the appearance of neutralizing antibodies against B.1.1.7 and B.1.351. The titers remained lower against B.1.351, even in responders. The vaccine displays a protective efficacy against COVID-19 as soon as two weeks after the first dose ^36^. Our results suggest that the low neutralizing titers (ED50 of 50-100) in the sera correspond to a protection against severe disease.

**Fig. 4.**
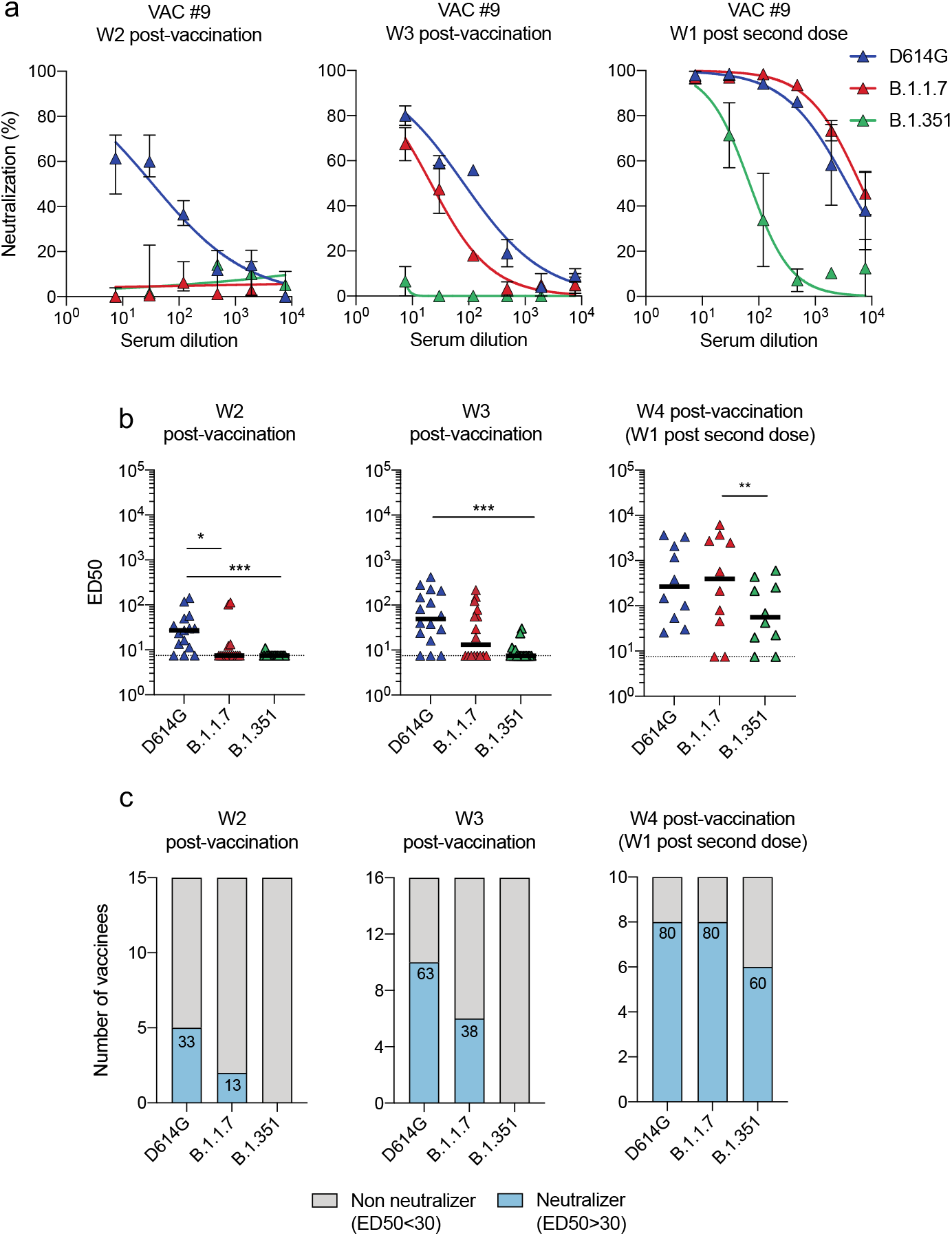
Sensitivity of SARS-CoV-2 D614G, B.1.1.7 and B.1.351 variants to sera from early vaccine recipients. **a.** Examples of neutralization curves with sera from one donor at weeks 2 (W2), 3 (W3) post vaccination and weeks 4 post-vaccination (W1 post second dose). Results are shown as mean ± SD from two to four independent experiments. **b.** ED50 of neutralization of the three viral isolates. Sera from 16 to 10 vaccine recipients were sampled at W2 (n=15), W3 (n=16) and W1 post second dose (n=10). Data are mean from two to four independent experiments. Friedman test with Dunn’s multiple comparison was performed between each viral strain at the different time points, *: p-value<0.05 **: p-value<0.01 **** p<0.001. **c.** Each subject was arbitrarily defined as “neutralizer” (in blue) if a neutralizing activity was detected at the 1/30 serum dilution or “non-neutralizer” (in grey) if no activity was detected. The numbers indicate the % of neutralizers.

**Fig. 5.**
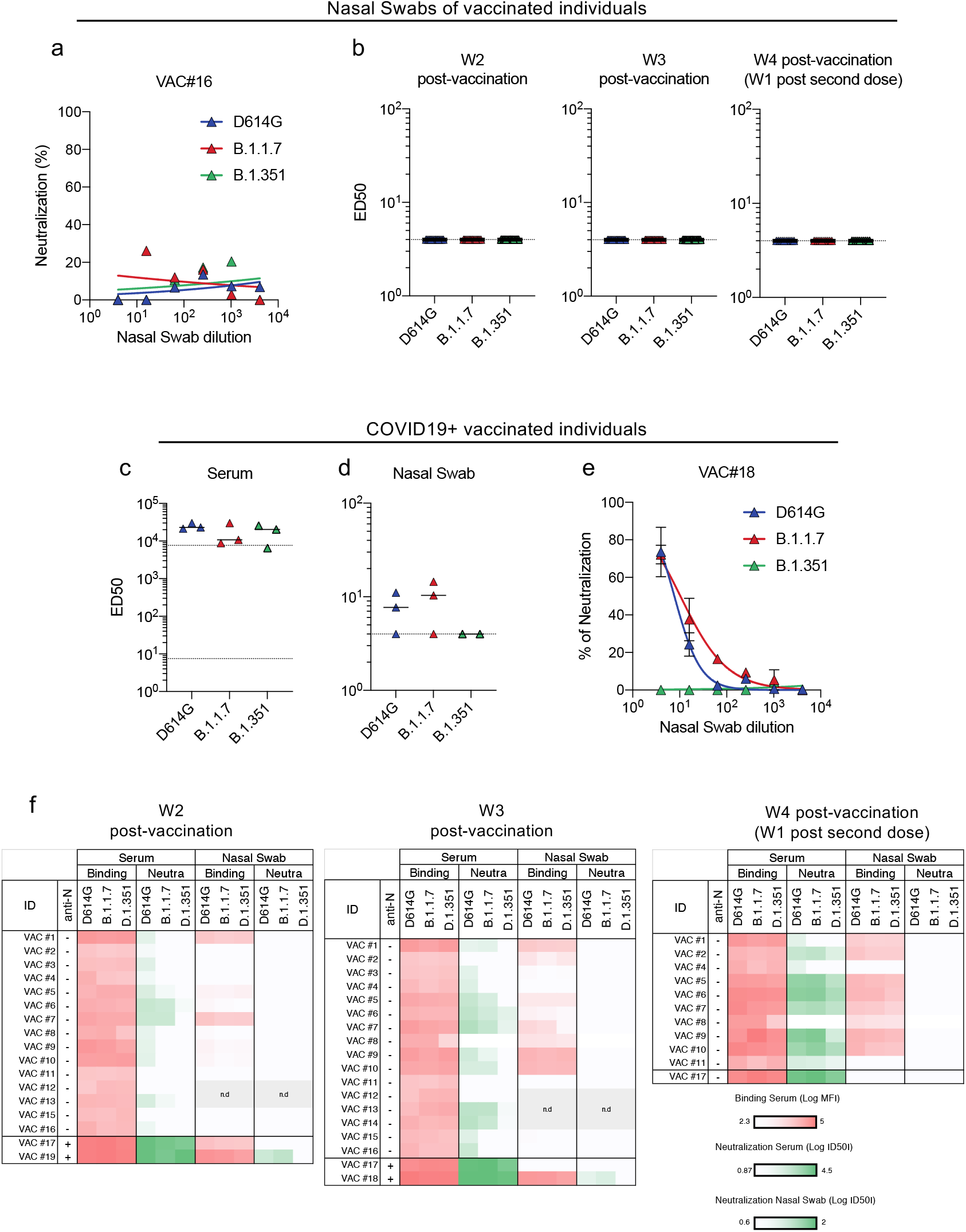
Sensitivity of SARS-CoV-2 D614G, B.1.1.7 and B.1.351 variants to nasal swabs from early vaccine recipients. **a.** Example of neutralization curves with nasal swab from one donor at week 2 (W2) post vaccination. No neutralization was detected. **b.** ED50 of neutralization of the three viral isolates, with nasal swabs from 16 vaccine recipients sampled at W2, W3 and W1 post-second dose. No neutralization was detected. Data are mean from two to four independent experiments. **c,d.** Analysis of the three COVID-19 vaccinated patients. Neutralizing antibody titers are shown in the serum and nasal swab, at week 2 post vaccination **e.** Example of neutralization curves with nasal swab from one such COVID-19 vaccine recipient. **f.** Heat map summarizing the levels of anti-Spike antibodies and their neutralizing activity in the sera and nasal swabs of the 19 vaccinated individuals. Vac #17, 18 and 19 were previously infected by SARS-CoV-2. White boxes denote an absence of activity.

### Sensitivity of SARS-CoV-2 B.1.1.7 and B.1.351 variants to nasal swabs from vaccine recipients

Little is known about the levels and function of vaccine-elicited antibodies in mucosal samples. We thus measured the neutralizing activity of nasal swabs from the series of vaccine recipients (Fig. 5). We did not detect any antiviral effect in these samples at two and three weeks post vaccination (Fig. 5a,b) We also analyzed three additional vaccine recipients who were seropositive for N at the time of vaccination, indicative of a previous infection. Two were diagnosed SARS-CoV-2 positive by PCR in March and April 2020 and experienced a mild disease. The third did not report any previous signs reminiscent of COVID-19. In these three individuals, the neutralizing titers were strikingly high at week 2 in their sera (ED50 of about 10^4^) (Fig. 5c). Two out of the three vaccine recipients who were previously seropositive for SARS-CoV-2 displayed a neutralizing activity in their nasal swabs (Fig. 5d,e). Their nasal swabs similarly neutralized the D614G and B.1.117 strains and were inactive against B.1.351. The levels of anti-Spike antibodies and their neutralization activity in the sera and nasal swabs of vaccine recipients are summarized in the heat map presented in Fig. 5f. In the swabs, we detected anti-SARS-CoV-2 antibodies in 6/17 available samples at week 2, 7/13 at week 3, 8/10 at week 4 (Fig. 5f). The three COVID-19+ vaccine recipients were those with highest antibody levels in the nasal swab. Unfortunately, we did not have access to their nasal swabs before vaccination, precluding determining whether previous infection, vaccination, or more likely the combination of both generated a neutralizing response in these individuals. Altogether, our results strongly suggest that vaccinees do not generally elicit an early humoral response detectable at mucosal surfaces. They strengthen the hypothesis that some vaccines may not protect against viral acquisition and infection of the oral-nasal region but may prevent severe disease associated with viral dissemination in the lower respiratory track.

## Discussion

Here, we analyzed the cross reactivity of the humoral response to pre-existing SARS-CoV-2 viruses and recently emerging variants, in sera from long-term convalescent individuals and recent vaccine recipients. We report that immune sera had slightly reduced but largely preserved activity against B.1.1.7 when compared to the reference D614G strain. The B.1.351 variant is more problematic as it is less sensitive or even unsensitive to a large part of the sera tested, particularly when global antibody levels are low. The E484K mutation, responsible for antibody escape of B.1.351 and P.1 variants, has been recently detected in the B.1.1.7 lineage in UK, and thus represents a major threat for previously immune populations. Here, we used authentic clinical viral isolates, rather than pseudovirus, providing a relevant way to assess inherent viral fitness and potential impact of additional mutations outside of Spike on sensitivity to neutralizing antibodies. On a technical note, the combination of S-Fuse and S-Flow assays, two methods of analysis of viral infectivity, neutralization or antibody levels, allowed a rapid assessment of the properties of emerging variants. The workflow established here can be easily adapted to any novel viral strain. Potential limitations of our study are the relatively low number of individuals analyzed and the short time frame of analysis after vaccination. Furthermore, we have not investigated the impact of pre-existing cellular immunity, which may be more cross-reactive against variants than the humoral response. Future work with more vaccine recipients that have or haven’t been previously infected, and longer periods of survey, will help characterizing the role of local and systemic humoral responses in vaccine efficacy. In conclusion, our results demonstrate that suboptimal or declining antibody responses are associated with a loss of cross-reactivity against novel emerging viral strains.

## Methods

No statistical methods were used to predetermine sample size. The experiments were not randomized and the investigators were not blinded to allocation during experiments and outcome assessment

### Orléans Cohort of convalescent and vaccinated individuals

Orleans’ Cohort of convalescent and/or vaccinated individuals: since April 2020, a prospective, monocentric, longitudinal, cohort clinical study enrolling 170 SARS-CoV-2-infected individuals and 30 non-infected healthy controls is on-going, aiming to describe the persistence of specific and neutralizing antibodies over a 24-months period. This study was approved by the ILE DE FRANCE IV ethical committee. At enrolment written informed consent was collected and participants completed a questionnaire which covered sociodemographic characteristics, virological findings (SARS-CoV-2 RT-PCR results including date of testing), clinical data (date of symptom onset, type of symptoms, hospitalization), and data related to anti-SARS-CoV-2 vaccination if ever (brand product, date of first and second vaccination). Serological status of participants was assessed every 3 months. Those who underwent anti-SARS-CoV-2 vaccination will have additional weekly blood and nasopharyngeal sampling after first dose of vaccine for a 2-months period (ClinicalTrials.gov Identifier: NCT04750720). For the present study, we randomly selected xx participants.

### Strasbourg Cohort of convalescent individuals

Since April 2020, a prospective, interventional, monocentric, longitudinal, cohort clinical study enrolling 308 RT-PCR-diagnosed SARS-CoV-2 infected hospital staff from the Strasbourg University Hospitals is on-going (ClinicalTrials.gov Identifier: NCT04441684). At enrolment (From April 15 to 29 2020) written informed consent was collected and participants completed a questionnaire which covered sociodemographic characteristics, virological findings (SARS-CoV-2 RT-PCR results including date of testing) and clinical data (date of symptom onset, type of symptoms, hospitalization). The serological status of the participants have been described at Months 3 (M3) and Months 6 (M6) POS ^33,34^. Laboratory identification of SARS-CoV-2 was performed at least 10 days before inclusion by RT-PCR testing on nasopharyngeal swab specimens according to current guidelines (Institut Pasteur, Paris, France; WHO technical guidance). The assay targets two regions of the viral RNA-dependent RNA polymerase (RdRp) gene with a threshold of detection of 10 copies per reaction. For the present study, we randomly selected 30 patients collected at M9.

### S-Fuse assay

U2OS-ACE2 GFP1-10 or GFP 11 cells, also termed S-Fuse cells, become GFP+ when they are productively infected by SARS-CoV-2 ^30^. Cells were tested negative for mycoplasma. Cells were mixed (ratio 1:1) and plated at 8×10^3^ per well in a μClear 96-well plate (Greiner Bio-One). The indicated SARS-CoV-2 strains were incubated with monoclonal antibodies, sera or nasal swabs at the indicated concentrations or dilutions for 15 minutes at room temperature and added to S-Fuse cells. 18 hours later, cells were fixed with 2% PFA, washed and stained with Hoechst (dilution 1:1,000, Invitrogen). Images were acquired with an Opera Phenix high content confocal microscope (PerkinElmer). The GFP area and the number of nuclei were quantified using the Harmony software (PerkinElmer). The percentage of neutralization was calculated using the number of syncytia as value with the following formula: 100 × (1 – (value with IgA/IgG – value in “non-infected”)/(value in “no IgA/IgG” – value in “non-infected”)). Neutralizing activity of each isotype was expressed as the half maximal effective concentration (IC50). IC50 values (in μg/ml for monoclonal antibodies and in dilution values for sera and nasal swabs) were calculated using a reconstructed curve using the percentage of the neutralization at the different indicated concentrations.

### Virus strains

The D614G strain (hCoV-19/France/GE1973/2020) was supplied by the National Reference Centre for Respiratory Viruses hosted by Institut Pasteur (Paris, France) and headed by Pr. S. van der Werf. This viral strain was supplied through the European Virus Archive goes Global (Evag) platform, a project that has received funding from the European Union’s Horizon 2020 research and innovation program under grant agreement n° 653316. The variant strains were isolated from nasal swabs on Vero cells and amplified by one or two passages on Vero cells. The B.1.1.7 strain originated from a patient in Tours (France) returning from United Kingdom. The B.1.1.351 strain (CNR 202100078) originated from a patient in Creteil (France). Both patients provided informed consent for the use of the biological materials. Titration of viral stocks was performed on Vero E6, with a limiting dilution technique allowing a calculation of TCID50, or on S-Fuse cells. Viruses were sequenced directly on nasal swabs, and after one or two passages on Vero cells. Sequences are available upon request.

### Spike expression plasmids

A codon optimized version of the SARS-Cov-2 Spike gene from Wuhan reference strain (GenBank: QHD43416.1), was transferred into the phCMV backbone (GenBank: AJ318514), by replacing the VSV-G gene. The B.1.1.7 (Δ69-70, ΔY144, N501Y, A570D, D614G, P681H, T716I, S982A, D1118H), and B.1.351 (L18F, D80A, D215G, Δ242-244, K417N, E484K, N501Y, D614G, A701V) Spike mutations were added in silico into the Wuhan codon optimized reference strain and were ordered as synthetic genes (GeneArt - Thermo Fisher Scientific) and transferred into the phCMV backbone. The D614G Spike was generated by introducing the corresponding mutation in the Wuhan reference strain using Q5® Site-Directed Mutagenesis Kit (NEB).

### Commercial serological assays

The serum samples from the Strasbourg cohort were first tested at Hôpital de Strasbourg using two commercial assays: 1) a CE-Marked LFA for detection of IgM and IgG against the SARS-CoV-2 RBD of the S protein developed by Biosynex® (COVID-19 BSS IgG/IgM). The test has a specificity of 99% and a sensitivity of 96% for samples >22 days POS. 2) a CE-Marked ELISA assay for detection of IgG against the full-length recombinant N protein from Epitope Diagnostics (EDITM Novel coronavirus COVID-19 IgG). The test has a specificity of 96% and a sensitivity of 81% after 28 days POS in our hands. The vaccine recipients were tested for anti-S antibodies at Hôpital Européen Georges Pompidou (Paris) with the following assays: Abbott SARS-CoV-2 IgG assays (Des Plaines, IL, USA) targeting SARS-CoV-2 nucleoprotein (N) were performed on Architect™ i2000SR analyzer (Abbott). Index value threshold for positivity was 1.4 as recommended. Beckman Coulter Access SARS-CoV-2 IgG assays (Brea, CA, USA) targeting the RBD were performed on UniCel DxI 800 Access Immunoassay System (Beckman Coulter). Index value threshold for positivity was 1 as recommended. Qualitative results as well as index values were used for analysis for both assays.

### S-Flow Assay

The S-Flow assay was performed as described ^33,35^. Briefly, HEK293T (referred as 293T) cells were acquired from ATCC (ATCC^®^ CRL-3216™) and tested negative for mycoplasma. 293T Cells were transfected with the indicated Spike expression plasmids or a control plasmid using Lipofectamine 2000 (Life technologies). One day after, transfected cells were detached using PBS-EDTA and transferred into U-bottom 96-well plates (50,000 cell/well). Cell were incubated at 4°C for 30 min with sera (1:300 dilution) or nasal swabs (1:50 dilution) in PBS containing 0.5% BSA and 2 mM EDTA, washed with PBS, and stained using anti-IgG AF647 (ThermoFisher), or Anti-IgM (PE by Jackson ImmunoResearch or AF488 by ThermoFisher). Cells were washed with PBS and fixed 10 min using 4% PFA. Data were acquired on an Attune Nxt instrument (Life Technologies). Stainings were also performed on control (293T Empty) cells. The specificity and sensitivity of this assay were originally assessed with the Wuhan Spike using 253 pre-pandemic samples and 377 RT-PCR confirmed SARS-CoV-2 cases. The sensitivity is 99.2% with a 95% confidence interval of 97.69%-99.78% and the specificity is 100% (98.5-100). Results were analysed with FlowJo 10.7.1 (Becton Dickinson). After testing 253 pre-pandemic samples, the positivity of a sample was defined as a specific binding above 40%. The specific binding was calculated as follow: 100 × (% binding 293T Spike ⍰ % binding 293T Empty)/ (100 - % binding 293T Empty).

### Antibodies and ACE-2 ectodomain

Human anti-SARS-CoV2 monoclonal antibodies were cloned from S-specific blood memory B cells of Covid19 convalescents (Planchais et al, manuscript in preparation). mAb48 and mAb102 recognize the RBD, and mAb10 binds to a conserved region of the S2 domain of the Spike protein. Human anti-S IgG mAbs and His-tagged recombinant ACE2 ectodomain (amino acids 19-615) cloned into pcDNA3.1 vector were produced by transient transfection of Freestyle^™^ 293-F cells and purified by affinity chromatography as previously described ^37^. Purified ACE2 protein was biotinylated using the EZ-Link Sulfo-NHS-Biotin kit (Thermo Fisher Scientific).

### Statistical analysis

Flow cytometry data were analyzed with FlowJo v10 software (TriStar). Calculations were performed using Excel 365 (Microsoft). Figures were drawn on Prism 9 (GraphPad Software). Statistical analysis was conducted using GraphPad Prism 9. Statistical significance between different groups was calculated using the tests indicated in each figure legend. All data associated with this study are available from O.S.

## Acknowledgments

We thank Felix Rey & Lisa Chakrabarti for critical reading of the manuscript. We thank patients who participated to this study, members of the Virus and Immunity Unit for discussions and help, Nathalie Aulner and the UtechS Photonic BioImaging (UPBI) core facility (Institut Pasteur), a member of the France BioImaging network, for image acquisition and analysis, Marija Backovic for the ACE2 construct. We thank the DRCI, CIC, Médecine du travail and Pôle de Biologie teams (CHU de Strasbourg) for the management of the Strasbourg cohort and serology testing.

**The authors do not declare any competing interest.**

## Author contributions

Experimental strategy design, experiments: DP, TB, LG, FGB, IS, FP, CP, JB, MMR, EB.

Vital materials MA, FD, SB, VE, MM, MG, JDS, HP, DV, AS, ESL, SFK, KS, HM, LH, SVDW, TP;

Manuscript writing: DP, TB, OS;

Manuscript editing: LD, JD, MMR, HP, DV, SFK, KS, HM, LH, SVDW, TP.

---

**Fig. S1.**
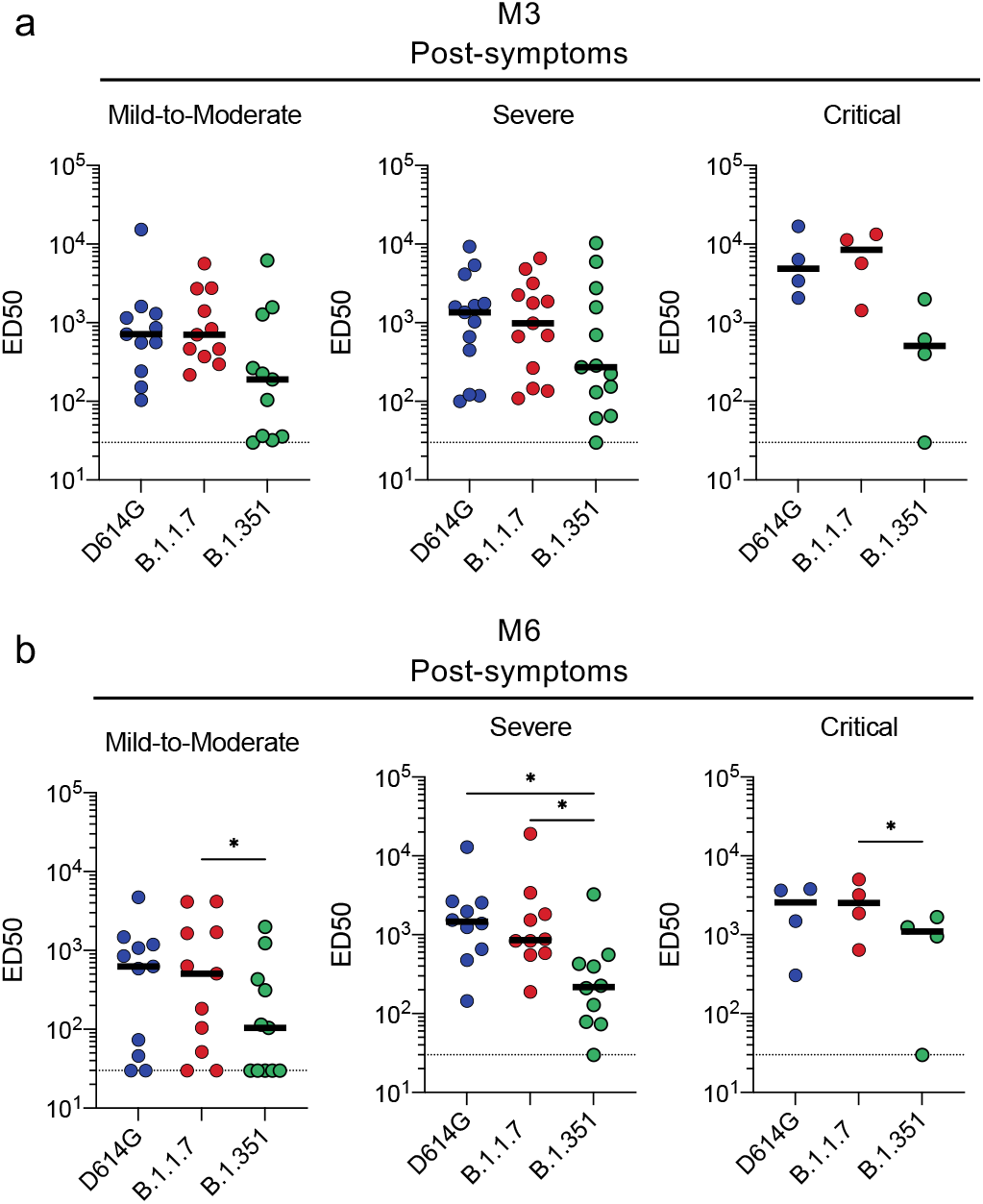
Sensitivity of SARS-CoV-2 variants to sera from different convalescent individuals. ED50 of neutralization of the D614G, B.1.1.7 and B.1.351. 32 sera from the Orléans Cohort were sequentially sampled at month 3 (M3) and M6 (upper and lower rows, respectively) and analyzed as described in Fig. 2 legend. The individuals are classified according to disease severity. Friedman test with Dunn’s multiple comparison was performed between each viral strain at the different time points, *: p-value<0.05 **: p-value<0.01 **** p<0.001.

**Fig. S2.**
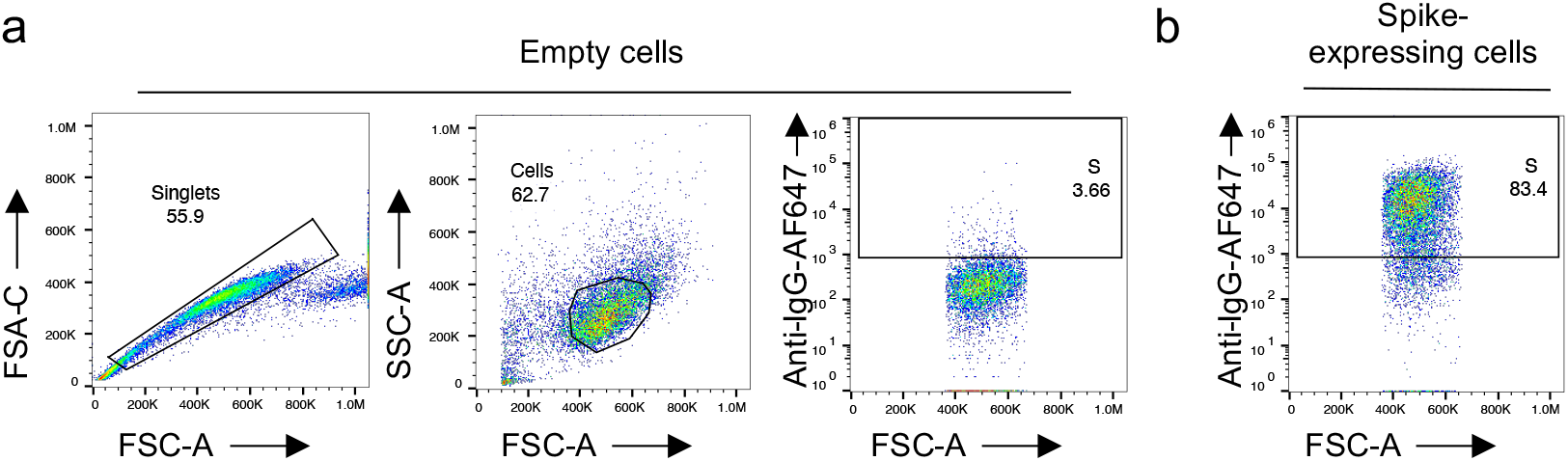
Gating strategy of the S-Flow assay. 293T cells were transiently transfected with plasmids expressing the D614G, B.1.1.7 and B.1.351 Spikes. After 24h, cells were stained with anti SARS-CoV-2 antibodies mAb10 (a pan-coronavirus antibody), mAb48, mAb102 or soluble ACE2 (ACE2-biotin at 10 μg/ml revealed with fluorescent Streptavidin) and analyzed by flow-cytometry. a. One representative example of the gating strategy is shown. Gates are set on cells transfected with an empty plasmid. b. An example of the signal obtained with a reactive serum on spike expressing cells is shown.

**Table. S1.** Characteristics of the 28 individuals from the Orléans cohort (upper panel) and 30 individuals from the Strasbourg cohort (upper panel).

| total n = 28 |  |  |
| --- | --- | --- |
| sex | female | 17 |
|  | male | 11 |
| Age | 56.9 (46-69) |  |
| Severity | Critical | 3 |
|  | Severe | 13 |
|  | Mild-Moderate | 11 |
|  | Asymptomatic | 1 |
| HIV | 2 |  |
| PCR | 26 |  |
| Anti-S (S-Flow) | 28 |  |
| 1st sampling (days) | 89 (71-105) |  |
| 2nd sampling (days) | 177 (164-197) |  |

| total n = 30 |  |  |
| --- | --- | --- |
| sex | female | 27 |
|  | male | 3 |
| Age | 44.8 (25-54) |  |
| Severity | Critical | 0 |
|  | Severe | 0 |
|  | Mild-Moderate | 30 |
|  | Asymptomatic | 0 |
| HIV | 0 |  |
| PCR | 30 |  |
| Anti-S (S-Flow) | 30 |  |
| Sampling (days) | 233 (206-258) |  |

**Table. S2.** Characteristics of the 19 vaccine recipients.

| total n = 19 |  |  |
| --- | --- | --- |
| sex | female | 6 |
|  | male | 13 |
| Age | 55 (46-63) |  |
| BMI | 24.2 (21.9-24.7) |  |
| Immune deficiency | 0 |  |
| Previous COVID-19 | 3 |  |
| Vaccine | Pfizer-BioNTech | 19 |
|  | Moderna | 0 |
|  | AstraZeneca | 0 |
| 1st shot | 4-13 january 2021 |  |
| Sampling | Week 2 | 17 |
|  | Week 3 | 18 |
|  | Week 4 | 11 |

